# GC-content evolution in bacterial genomes: the biased gene conversion hypothesis expands

**DOI:** 10.1101/011023

**Authors:** Florent Lassalle, Séverine Périan, Thomas Bataillon, Xavier Nesme, Laurent Duret, Vincent Daubin

## Abstract

The characterization of functional elements in genomes relies on the identification of the footprints of natural selection. In this quest, taking into account neutral evolutionary processes such as mutation and genetic drift is crucial because these forces can generate patterns that may obscure or mimic signatures of selection. In mammals, and probably in many eukaryotes, another such confounding factor called GC-Biased Gene Conversion (gBGC) has been documented. This mechanism generates patterns identical to what is expected under selection for higher GC-content, specifically in highly recombining genomic regions. Recent results have suggested that a mysterious selective force favouring higher GC-content exists in Bacteria but the possibility that it could be gBGC has been excluded. Here, we show that gBGC is probably at work in most if not all bacterial species. First we find a consistent positive relationship between the GC-content of a gene and evidence of intra-genic recombination throughout a broad spectrum of bacterial clades. Second, we show that the evolutionary force responsible for this pattern is acting independently from selection on codon usage, and could potentially interfere with selection in favor of optimal AU-ending codons. A comparison with data from human populations shows that the intensity of gBGC in Bacteria is comparable to what has been reported in mammals. We propose that gBGC is not restricted to sexual Eukaryotes but also widespread among Bacteria and could therefore be an ancestral feature of cellular organisms. We argue that if gBGC occurs in bacteria, it can account for previously unexplained observations, such as the apparent non-equilibrium of base substitution patterns and the heterogeneity of gene composition within bacterial genomes. Because gBGC produces patterns similar to positive selection, it is essential to take this process into account when studying the evolutionary forces at work in bacterial genomes.

## Introduction

Comparative genomics is a fundamental key to the inner workings of genomes. The identification of genes and other functional elements such as regulatory regions, as well as the understanding of their influence on the fitness of organisms rely essentially on the detection of signatures of natural selection within genomes [1]. In that respect, devising a model of sequence evolution in the absence of selective constraints (a neutral model) is critical for the detection of functional sequences. Indeed, to explain the features of a given genomic segment, comparing the fit of a neutral model to that of a model that also invokes selection (either purifying or positive) is the operational way to infer evolutionary constraint and hence function.

The base composition of genomic sequences varies widely, both across species and along chromosomes [2,3]. For instance, the genomic GC-content of cellular organisms ranges from 13% to about 75% [4,5], with vast intra-genomic heterogeneity. These large-scale variations in base composition affect all parts of genomes, intergenic regions and genes – including all three codon positions [6] – and hence cannot be simply explained by selective constraints on the encoded proteins. Determining the underlying causes (selective or neutral) of these variations in GC-content is a major issue in genetics: if they result from selection, it implies that the genomic base composition *per se* is an important trait that contributes to the fitness of organisms; conversely, if these “genomic landscapes” are largely shaped by non-adaptive molecular processes, then characterizing these processes is essential for the reliable detection of selection (see e.g. [7]).

In mammals, the analysis of polymorphism data and substitution patterns along genomes demonstrated that the evolution of GC-content is driven by recombination, which tends to increase the probability of fixation of AT→GC mutations [8,9]. The impact of recombination on base composition in these genomes is most probably due to a phenomenon known as GC-biased gene conversion (gBGC), which favours G/C nucleotides at polymorphic sites in the conversion of intermediates of recombination (see review in [10]). Although gBGC as a process is unrelated to natural selection, it affects the probability of fixation of alleles in patterns similar to selection [11]. It has been shown to be an important confounding factor, which can mimic some marks of positive selection [7,12] and interfere with selection by actively promoting the fixation of deleterious alleles [13,14]. The process of gBGC has been observed directly in meiosis products from yeast and human [15,16], and there is ample evidence, based on the analysis of relationships between recombination rate and substitution patterns within genomes, that this process affects many other eukaryotes [17–19].

In Bacteria and Archaea, several environmental factors potentially affecting genomic GC-content have been proposed (such as the availability of oxygen or nitrogen in the environment, growth temperature, or the variety of environments encountered by an organism, see for instance [20] and ref. therein). Because these effects are weak and the nature of the selective pressures remain elusive, the major force driving genomic GC-content has long been considered to be mutational bias [21]. Recently however, two independent analyses have shown that in virtually all Bacteria, independently of their genomic GC-content, there is an excess of G/C→A/T mutations [22,23]. This suggests that an unknown process, selective or neutral, is opposing this universal mutational bias by favouring the fixation of G/C alleles Previously, an analysis of a large number of *E. coli* genomes had suggested a possible role of gBGC, based on the link between GC-content, recombination and the organization of the chromosome in this species [24]. However Hildebrand et al. [23] observed that the excess of G/C→A/T mutations was still present after removing datasets with evidence of recombination. Moreover they found no correlation between GC-content and recombination rate across bacterial species. They therefore concluded that this force could not be gBGC and hence that selection was driving an increase of genomic GC in Bacteria. The nature of this selective advantage remains however mysterious, though various hypotheses have been proposed [25,26].

Here we argue that the analyses performed by Hildebrand et al. [23] are not conclusive regarding the gBGC hypothesis, and we present evidence that variations in GC-content observed in Bacteria are influenced by gBGC. One pervasive signature of gBGC is that genomic regions undergoing high recombination rates will also acquire a high GC-content [6]. We thus studied the relationship between recombination and GC-content in 20 groups of Bacteria and one group of Archaea. This dataset covers a wide range of clades representative of the bacterial diversity. To avoid problems inherent to comparisons of recombination rates among species (such as differences in polymorphism, genome samples, population size, mutation rates, an other life history factors), we examined the intragenomic variability for both recombination and GC-content.

We show that in a wide variety of bacterial species, genes with evidence of recombination have a higher GC-content. We further show that this bias towards G/C nucleotides in recombining genes cannot be explained by selection on codon usage, and could interfere with the selection for AT-ending optimal codons. These two observations strongly suggest that homologous recombination, *via* gBGC, is a crucial factor universally influencing the nucleotide content of genes and genomes. If confirmed, gBGC can account for several pervasive yet unexplained features of bacterial genomes. Finally, we emphasize that because gBGC has the ability to both mimic and interfere with natural selection, gBGC must be considered by future studies geared at understanding processes driving bacterial genome evolution.

## Results

### A universal relationship between recombination and GC% in Bacteria

In Bacteria, recombination occurs in the form of gene conversion (i.e. unidirectional transfer of genetic material from a donor sequence towards a homologous recipient sequence). To detect past gene conversion events in bacterial species, it is necessary to compare closely related genomes. We therefore selected in the database of homologous gene families HOGENOM (release 6) [27] all groups of closely related species or strains encompassing at least 6 sequenced genomes. This dataset contains 20 bacterial groups and one archaeal group. For each gene family represented in these groups, we computed i) the average GC-content at different positions of codons and ii) the index of recombination provided by PHI [28] based on alignments of standardized length (see methods for details). PHI is a rapid method for detecting recombination in multiple alignments at the scale of the gene, which has been shown to be more robust than most methods to variations in recombination rates, sequence divergence and population dynamics [28]. We used this test to determine if homologous gene families had experienced gene conversion events among members of the taxa of interest. One important feature of this test is that it measures whether there is sufficient phylogenetic signal in an alignment to tell if recombination has occurred. Only those alignments with sufficient signal, whether recombinant or non-recombinant were retained for tests in the remaining of this study. We also used three other approaches for detecting recombination, and these confirm the robustness of our conclusions (see Methods and Supplementary Material).

In Eukaryotes, a general relationship between various estimates of recombination rate and the GC% of genes has been documented and provides indirect evidence for gBGC. Our first goal was to test this prediction in Bacteria and Archaea. To exclude a potential effect of the number of genes in the alignment on our estimates of recombination (because alignments with more sequences are expected to give more power to detect recombination), we focused on single-copy genes of the core genome (i.e. genes that are present in only one copy and found in each genome of a group). In 7 of the 21 groups, the proportion of single-copy genes of the core genome with evidence for recombination was very low (<2% of all gene alignments tested), suggesting that these species are clonal or nearly so (Table 1; shaded datasets in Fig. 1). In 11 of the 14 remaining groups, we found a significant positive difference in average GC-content at all and/or at the third position of codons (GC3) between recombinant and non-recombinant genes (Figure 1). In these 11 species, the difference in GC3 is always larger than that at all positions, suggesting that the effect of recombination on gene composition is stronger at synonymous positions (probably because of purifying selection on protein sequences). Two notable exception to this pattern are i) the bacterial species *Helicobacter pylori*, where GC-content seems to be lower in recombining genes and ii) the *Bacillus anthracis/cereus* group, where GC at all positions and GC3 display opposite patterns, with GC3 being higher in recombining genes. Consistent results are obtained using alternative recombination detection methods (Fig. S1).

**Table 1:** Dataset used in this study. Total number of genes in the core-genome, as well as the number of core genes classified as recombinant and non-recombinant based on PHI analysis and unclassified ones (genes with insufficient signal to test for recombination, excluded from comparison tests) are indicated. The mean proportion of GC (all positions of codons) and GC3 (third position of codons) of core genes are shown for each dataset.

| Dataset | Taxon name | Nb. of<br>genomes | Total nb.<br>of core<br>genes | Nb. of core<br>genes of<br>size ≥<br>900bp | Nb. of<br>recombinant<br>core genes | Nb. of non-recombinant<br>core genes | Nb. of<br>unclassified<br>core genes* | Mean GC<br>of core<br>genes (%) | Mean<br>GC3 of<br>core<br>genes (%) | Mean GC of<br>core genes of<br>size ≥ 900bp<br>(%) | Mean GC3 of<br>core genes of<br>size ≥ 900bp<br>(%) |
| --- | --- | --- | --- | --- | --- | --- | --- | --- | --- | --- | --- |
| Brsp | *Brucella* spp. | 9 | 1675 | 776 | 0 | 422 | 354 | 58,1 | 67,3 | 58,8 | 68,6 |
| Ftul | *Francisella tularensis* | 8 | 1015 | 469 | 0 | 359 | 110 | 33,3 | 21,8 | 33,8 | 21,9 |
| Mtub | *Mycobacterium tuberculosis* complex | 7 | 2222 | 1078 | 0 | 91 | 987 | 65,6 | 79,3 | 66,1 | 80,3 |
| Bmal | *Burkholderia Pseudomallei* group | 9 | 1482 | 781 | 4 | 628 | 149 | 67,9 | 88,3 | 68,7 | 89,7 |
| Ypes | *Yersinia pestis* | 11 | 2017 | 1073 | 7 | 679 | 387 | 48,7 | 48,9 | 49,3 | 49,9 |
| Ctra | *Chlamydia trachomatis* | 13 | 772 | 391 | 6 | 376 | 9 | 41,6 | 34,6 | 41,8 | 34,6 |
| Susp | *Sulfolobus* spp. | 8 | 1386 | 547 | 8 | 465 | 74 | 34,7 | 29,1 | 35,4 | 29,2 |
| Bcen | *Burkholderia cenocepacia* complex (BCC) | 8 | 1939 | 936 | 106 | 829 | 1 | 67,4 | 87,4 | 68,2 | 89,1 |
| Saur | *Staphylococcus aureus* | 15 | 1464 | 668 | 92 | 571 | 5 | 33,5 | 22,4 | 34,2 | 22,2 |
| Blon | *Bifidobacterium longum* | 6 | 1006 | 600 | 73 | 358 | 169 | 61,3 | 77,2 | 61,9 | 78,2 |
| Abau | *Acinetobacter* spp. | 6 | 1429 | 638 | 115 | 516 | 7 | 40,3 | 29,9 | 40,8 | 30,2 |
| Cjej | *Campylobacter jejunii* | 6 | 1048 | 501 | 93 | 403 | 5 | 31,0 | 19,4 | 31,6 | 19,6 |
| Cbot | *Clostridium botulinum* | 8 | 1715 | 730 | 168 | 562 | 0 | 28,6 | 16,8 | 29,1 | 16,4 |
| Spyo | *Streptococcus pyogenes* | 12 | 1051 | 496 | 119 | 370 | 7 | 38,9 | 31,6 | 39,6 | 32,2 |
| Spne | *Streptococcus pneumoniae* | 13 | 1090 | 507 | 119 | 368 | 20 | 41,3 | 36,9 | 42,0 | 37,4 |
| Sent | *Salmonella enterica* | 14 | 2121 | 975 | 323 | 652 | 0 | 53,4 | 59,4 | 54,6 | 61,6 |
| Lisp | *Listeria* spp. | 8 | 1631 | 678 | 328 | 350 | 0 | 38,1 | 29,6 | 38,8 | 29,5 |
| Bant | *Bacillus anthracis/cereus* group | 17 | 1730 | 727 | 422 | 305 | 0 | 36,3 | 26,3 | 37,0 | 26,5 |
| Nmen | *Nesseiria meningitidis* | 8 | 1156 | 552 | 339 | 213 | 0 | 54,4 | 63,9 | 55,3 | 65,7 |
| Ecol | *Escherichia coli* | 35 | 1357 | 619 | 442 | 177 | 0 | 52,3 | 56,3 | 53,3 | 58,0 |
| Hpyl | *Helicobacter pylori* | 14 | 995 | 467 | 334 | 133 | 0 | 39,9 | 42,8 | 40,4 | 43,3 |

**Figure 1.**
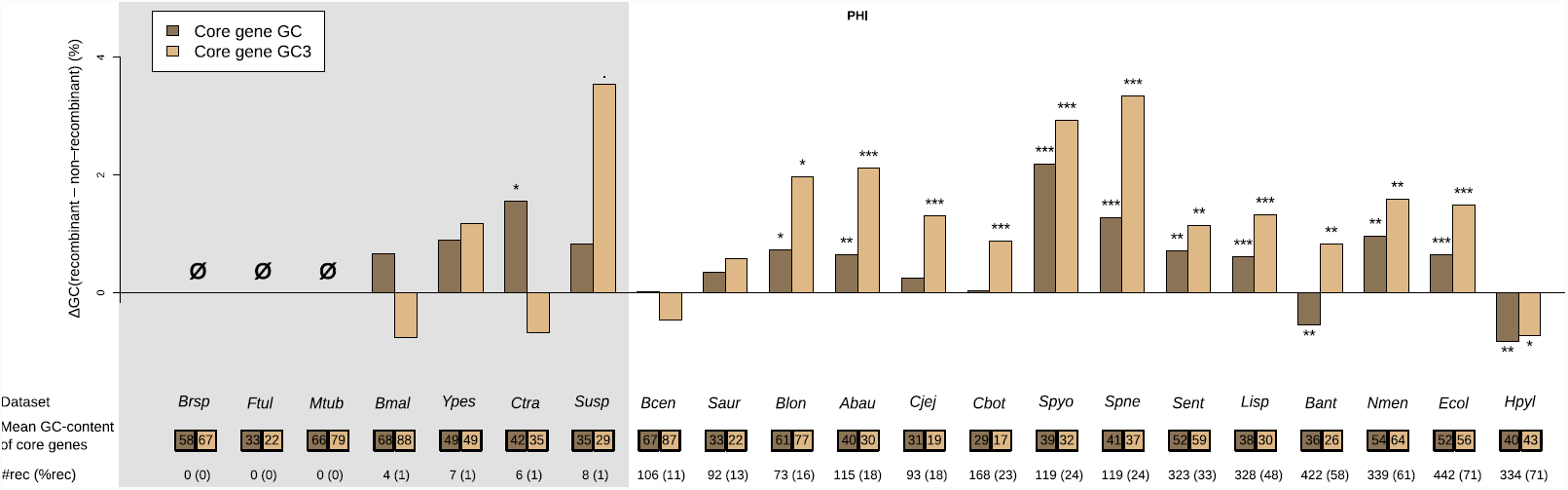
Effect of recombination on core genes GC-content. Difference in average GC-content of recombinant and non-recombinant genes, measured on entire CDS length (GC, dark brown) or at third codon position only (GC3, light brown) in the core genome of each dataset. Recombinant status was determined using the PHI test [28] (*p* < 0.05) on alignments of core gene families trimmed to a common length of 900bp. A positive difference indicates that recombinant families are enriched in GC. Stars indicate the level of significance of a Student’s *t*-test (“.”, *p* < 0.1; “*”, *p* < 0.05; “**”, *p* < 0.01; “***”, *p* < 0.001). Statistical tests are detailed further in Table S1. Dataset abbreviations are explained in Table 1. Boxes under dataset names indicate the mean percentage of GC and GC3 values of core genes. Figures at the bottom indicate the number of recombinant core gene families in the dataset, figures in parenthesis indicate their percentage in the total pool of recombinant and non-recombinant families. Shading in background marks datasets with less than 20 recombinant gene families (detailed Table 1).

In principle, gBGC should affect all genomic regions where recombination occurs, including intergenic regions. Intergenes are generally shorter than coding regions. Furthermore, they evolve more rapidly and hence are more difficult to align. Hence, the methods that we used to detect recombination in coding regions cannot be applied with intergenes. We therefore used the recombinant or non-recombinant status of the neighboring protein-coding genes as a proxy of the status of the intergenes. In 11 of the 14 taxonomic groups, we observed that intergenes flanked by recombining genes have a higher GC-content than intergenes flanked by non-recombining genes (Fig. S2). The difference in GC-content between the two classes is weaker than that observed in coding regions, and when considered individually, only one comparison is statistically significant (*Streptococcus pyogenes*, *p* < 0.01). This is possibly because the prediction of recombination status of intergenic regions is indirect (based on the status of flanking genes) and therefore less accurate than that of coding regions. However, the number of cases where the difference in GC-content between the two classes is positive (11 out of 14 comparisons) is significantly higher than expected by chance (Chi-squared test: p=0.03).

### Selection on optimal codons cannot explain the association of GC with recombination

Recombination is known to enhance the efficacy of selection by breaking linkage between neighboring selected sites. It is therefore possible that selection is more efficient in recombining genes. This effect (referred to as Hill-Robertson interference) should theoretically be more pronounced in the case of selection on codon usage [29], which is relatively weak compared to selection on amino acid sequences. Thus recombination – in the absence of any gBGC – can potentially explain the pronounced effect observed on GC3: if Hill-Robertson interference leads to a higher frequency of optimal codons in highly recombining genes, and if optimal codons tended to be GC-rich, this effect could explain a relationship between GC-content and recombination. The “selection model” sketched above predicts that the frequency of all optimal codons (both GC-ending and AU-ending) should increase with recombination. In contrast, a model incorporating the effect of gBGC predicts that GC-ending codons (not specifically optimal codons) should be enriched in recombining regions, and that AU-ending codons (and possibly AU-ending optimal codons if gBGC is strong enough to override selection on codon usage) should display the opposite pattern. We therefore looked specifically at the frequency of the different types of codons, i.e. optimal and non-optimal, in recombining and non-recombining genes. There is a debate over the best way to define optimal codons, based on their over-representation in either ribosomal protein genes (RP), or genes with the highest codon bias (HCB) [30–32]. We therefore analyzed the frequency of GC-ending and AU-ending optimal codons (FopGC and FopAU) and non-optimal codons (FnopGC and FnopAU) according to both RP and HCB definitions (Fig. 2 and S3 and Fig. S4, respectively). The higher GC3 of recombining genes means that GC-ending codons are over-represented in recombining genes, but this is true for optimal GC-ending codons in only 2 (RP optimal codons) or 4 (HCB optimal codons) species out of 11. This effect is hence essentially due to non-optimal codons (FnopGC is significantly higher in recombining genes than non-recombining genes in respectively 9 and 8 species for RP and HCB definitions). Moreover, optimal AU-ending codons are significantly depleted in recombining genes for 8 (resp. 5) species for RP (resp. HCB) codons. In fact, only two species, *S. pyogenes* and *Nesseiria meningitidis* (using the HCB method – only *S. pyogenes* using the RP method) exhibit a pattern partially compatible with the selection hypothesis presented above. All species display either an increase of FnopGC and/or a decrease of FopAU in recombining genes, a fact that cannot be explained by a higher efficiency of selection. This pattern excludes the possibility of pervasive selection for codon usage promoting a better adaptation to the pool of tRNA for genes in regions of high recombination, but is compatible with the predictions of gBGC.

**Figure 2.**
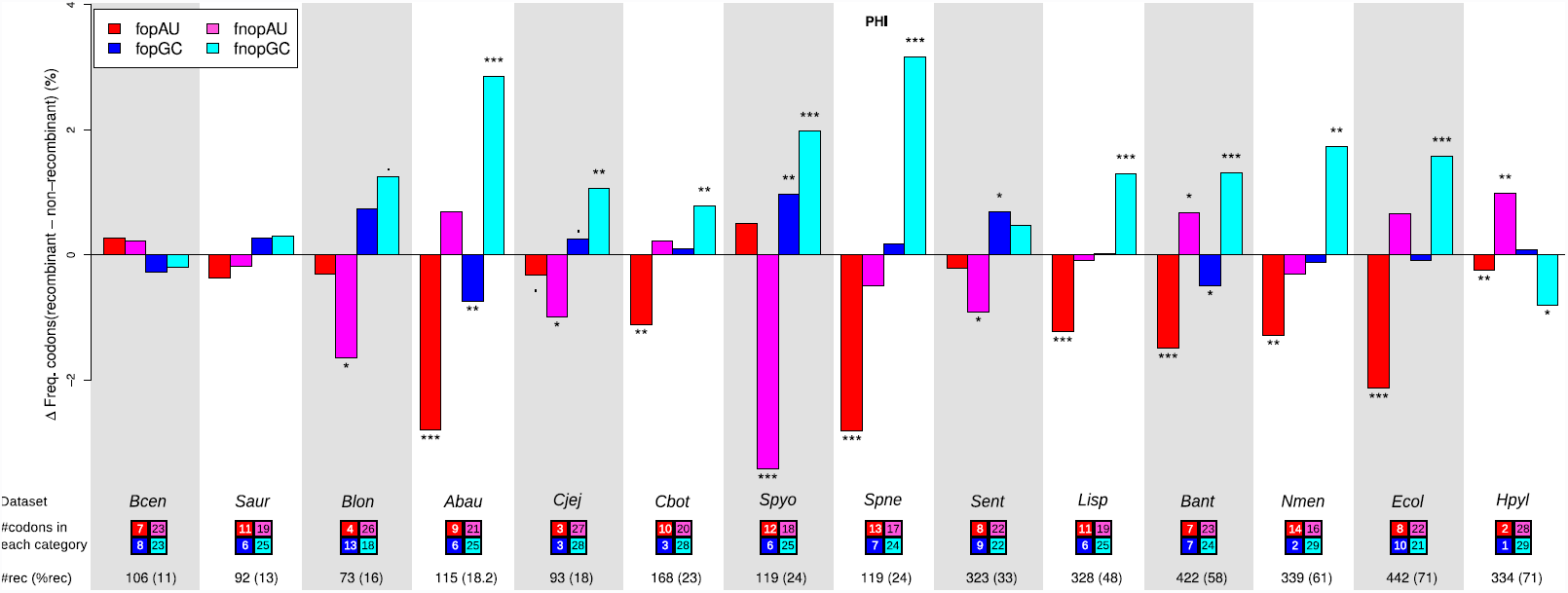
Effect of recombination on codon usage of core genes. Difference in frequency of optimal (fop) or non-optimal (fnop) codons (as determined by RP method) in recombining and non-recombining genes in each dataset for AU-ending (redish colors) and GC-ending (blueish colors) codons. The recombination status of genes was determined as in Fig. 1, only datasets with more than 10% recombining genes are shown. A positive difference indicates that recombining genes are enriched in a category of codons, while a negative difference indicate depletion. Stars indicate significance of a Student’s *t*-test between recombining and non recombining genes. Boxes under dataset names indicate the numbers of AU-ending and GC-ending optimal or non-optimal codons used by the taxon (detailed in Table S2). Symbols and dataset abbreviations as in Fig. 1; shading is only used to distinguish between datasets. It should be noticed that variations in fopGC and fnopAU (resp. fopAU and fnopGC) are not totally independent (typically, for all amino-acids encoded by two synonymous codons, if the optimal codon is GC ending, the non-optimal is AT-ending).

### Quantifying the impact of recombination on bacterial genome evolution

To quantify the relationship between recombination and base composition, we first analyzed the genome of *S. pyogenes*, one of the species for which the signature of gBGC is strong (Fig. 1, see also Sup. Mat.). We used ClonalOrigin [33] to compute the population-scaled recombination rate (*rho*) for each gene of the core genome. The correlation between *rho* and the GC-content at third codon position of each gene (GC3) is slight but significant (R^2^=0.034, p<10^-4^, n=478). Interestingly, when we exclude genes for which the estimate of *rho* is less reliable (about 10% of the data, see Sup. Mat.), the correlation strongly increases (R^2^=0.087; p<10^-9^, n=437). Due to the low amount of data available in a gene-scale alignment, the measure of *rho* is expected to be noisy. Thus, the observed correlation is probably an underestimate. To try to get more robust estimates of *rho*, we binned the dataset into 20 groups of genes according to their GC3, and we computed the correlation between the average GC3 and the average *rho* of each bin. Using this approach, we observed a strong correlation between the GC3 and recombination (R^2^=0.60; Fig. 3A).

**Figure 3.**
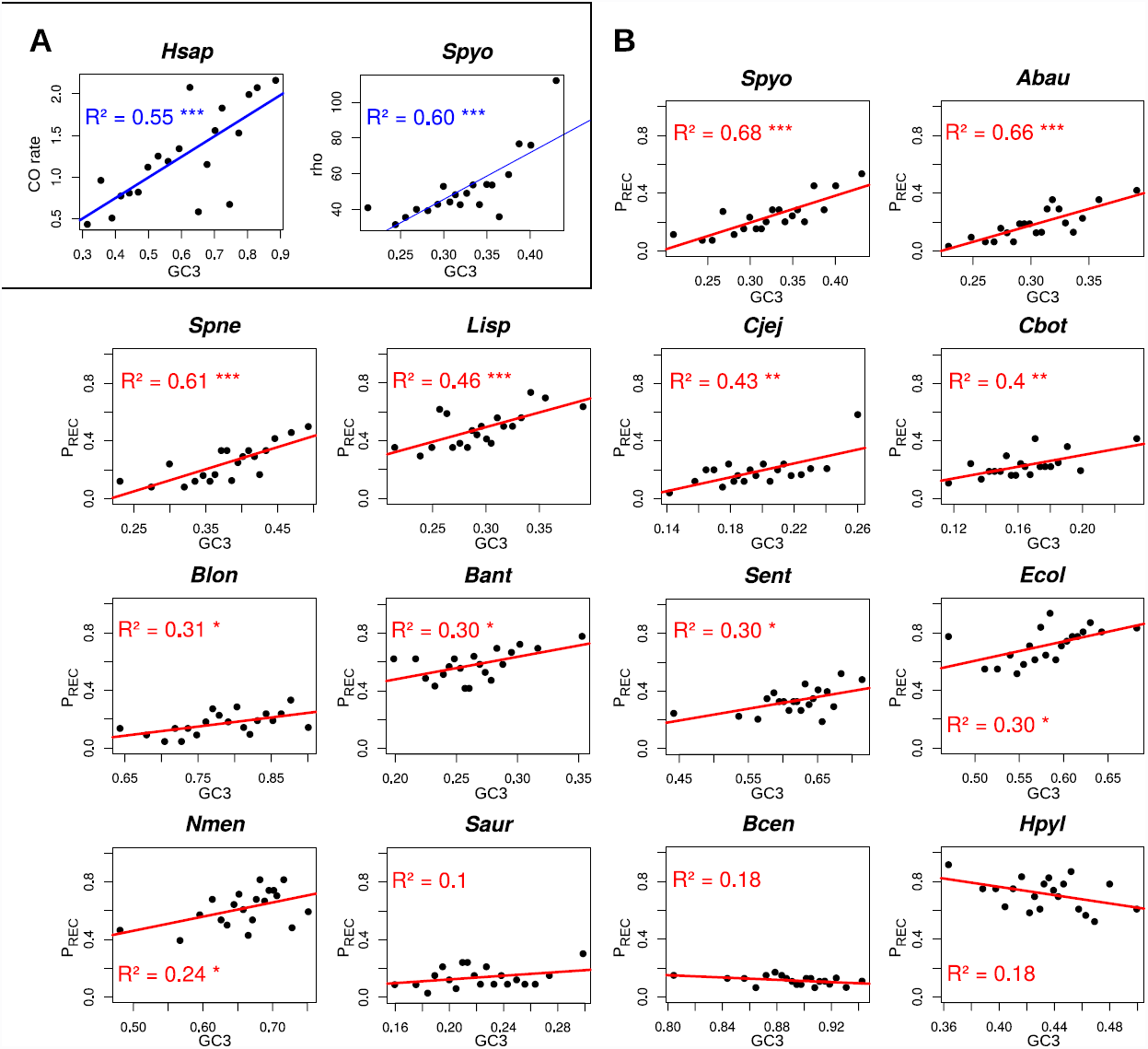
Correlations between GC3 and estimates of recombination rate. For each dataset, core genes are sorted by increasing GC3 and pooled into 20 classes of equal size. Correlations between the mean GC3 and mean recombination rate of each class are reported. (A) Correlation between GC3 and coalescent-based estimates of recombination rate for *Homo sapiens* (*Hsap*) and *Stretococcus pyogenes* (*Spyo*). For *Hsap*, recombination rate is expressed as cM·^Mb-^1; a subset of 600 genes out of the 16,346 human genes is shown as a representative of 1,000 random samples (mean R^2^ is 55%, see Main Text). For *Spyo*, recombination rate is expressed as the value of *rho* parameter in ClonalOrigin [33] inferences, which is scaled by arbitrary coalescent time units; a subset of 437 genes out of 478 core genes was used, after removal of the 41 genes showing no convergence of the *rho* estimate (correlation on the full 478 core genes yields a R^2^ of 31%, see Suplementary Text). (B) Correlation between GC3 and P_REC_, the proportion of genes detected as recombinant by PHI test [28] in the class, for all 14 bacterial datasets showing sufficient evidence of recombination (Table 1).

For a comparison, we performed a similar analysis in humans: we randomly selected 600 human genes (which corresponds to the average number of genes analyzed in our bacterial data sets), binned the data set into 20 groups of genes according to their GC3, and we computed the correlation between the average GC3 and the average recombination rate as obtained from population-wide surveys [34]. The average correlation (computed after repeating the random sampling 1,000 times) is R^2^=0.55 (with 95% of the R^2^ values in the interval [0.26-0.78]; one representative example is presented in Fig 3A).

Using this binning approach, we noted that the proportion of genes in a bin that are detected as recombinant by PHI is strongly correlated with the average value of *rho* (R^2^=0.70; p<10^-5^). This suggests that this index (hereafter referred to as P_REC_, for ‘proportion of recombinant’) is a good proxy for the average recombination rate in a bin. In fact, we observed that GC3 correlates more strongly with P_REC_ (R^2^=0.68; Fig. 3B) than with *rho*. Given that the computation of *rho* with ClonalOrigin is extremely time consuming, we decided to use P_REC_ to evaluate the correlation between GC3 and recombination rate in other species. Correlations were positive and significant for 11 of the 14 species, with R^2^ values ranging from 0.24 to 0.68, and we did not observe any significant negative correlation (Fig. 3B). This shows that the correlations between GC-content and recombination rate in bacteria are of similar magnitude to what is observed in humans.

## Discussion

### Is there selection for higher genomic GC-content in bacteria?

Our results suggest that recombination affects the GC-content of genes in most bacterial phyla. We analyzed genes of the core genome to compare the base composition of genes with or without evidence of recombination. In our sample, seven bacterial species showed very little evidence of recombination (less than 2% of gene alignments with detectable traces of recombination): the *Burkholderia pseudomalei* group, *Chlamydia trachomatis, Francisella tularensis, Mycobacterium tuberculosis* and *Yersinia pestis* which are species known to be pathogenic clonal complexes with low polymorphism and probably very low recombination [35–39], while *Brucella spp.* and *Sulfolobus spp.* are likely composed of ecologically isolated clades, because of their respective lifestyle as obligate intracellular pathogen or ecotypes endemic of hot springs [40–42]. The 14 other bacterial species contain clear signal of recombination (11% to 71% of testable core genes with evidence of recombination). In 11 of these 14 species, we observed that the GC-content (measured at the third codon position or along the entire coding region) is higher among recombining genes compared to other (hereafter labeled as “non-recombining”) core genes.

Several hypotheses have been proposed to explain the variations in GC-content among bacterial genomes [25]. Recently, two studies have revealed that the genomic GC-content of bacterial genomes is always higher than what would be predicted from mutational bias [22,23]. Hence, it seems inescapable that some other evolutionary force is driving the genomic GC-content towards higher values in virtually all bacterial species, except maybe for the most AT-rich genomes [22]. A former study of *E.coli* genomes showed that the excess of new AT-enriching mutation was erased with time, but that purifying selection on protein sequence could not alone account for this compensatory process [43]. Actually, the recombination-associated process we observe here is noticeably stronger at the third codon position, which is often synonymous. This suggests that purifying selection on protein sequences is rather counteracting its effect. Recombination is known to enhance the efficiency of selection by breaking linkage among sites. It is therefore conceivable that our results merely reveal a universal selective pressure favoring GC-rich alleles. But the mechanism underlying such selection would have to be acting more efficiently on synonymous sites than non-synonymous sites because the difference of GC% between recombining and non-recombining genes is higher at the third position of codons. This excludes potential selection on amino-acid content. One selectable trait that may influence synonymous positions is codon usage. If optimal codons tended to be GC-rich, recombination could drive GC% higher by favoring the adaptation of genes to better translation efficiency. However, we observed a higher GC-content in recombining genes even in species favoring A/U-ending codons (Fig. 2). Moreover, A/U-ending and G/C-ending codons show opposite relationships with recombination in most species, irrespective of their optimality. These observations suggest that the evolutionary force explaining our results is also largely independent from selection on codon usage. This conclusion is supported by the fact that the relationship between GC-content and recombination is also observed in intergenic regions (Fig. S2).

In fact, as suggested by Hershberg and Petrov [22], who observed that the intergenic regions of bacterial genomes also have higher GC% than expected from their mutational pattern, it seems likely that the process is unlinked to gene expression or function. Hence, either there is selection acting simultaneously on each nucleotide of a bacterial genome to become G or C, or GC-biased gene conversion, which has now been observed in a variety of Eukaryotes is also at work in Bacteria.

### gBGC effect on nucleotide composition is manifest within genomes

The hypothesis that gBGC plays a role in bacterial genome evolution has been considered previously [23]. Hildebrand et al. analyzed the correlation between genome-wide measures of recombination rate (scaled by effective population size) with genomic GC-content among 34 species, covering different bacterial phyla. As they did not find any significant correlation, they concluded that there was no evidence of gBGC in Bacteria [23]. However, we argue here that this observation is not conclusive. In fact, the strength of gBGC depends on four variables: the effective population size (*N_e_*), the rate of recombination per bp per generation (*r*), the length of conversion tracts (*L*) and the intensity of the repair bias (*b_0_)* (for review, see [6]). In a haploid organism, the population-scaled gBGC coefficient is:

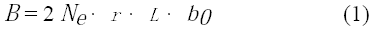

Similar to selection, the impact of gBGC on genome evolution depends on its intensity relative to genetic drift, and becomes negligible when *B* ≪ 1. There is evidence that besides *N_e_* and *r*, both *L* and *b_0_* can vary strongly across species. For example, in budding yeast, when a GC/AT heterozygote site is involved in a gene conversion event, the GC-allele is transmitted with a probability *p_GC_*=0.507 (which is significantly higher than the expected Mendelian transmission ratio; [15]), whereas in humans, a recent analysis of gene conversions tracts associated to non-crossover recombination showed that GC-alleles are transmitted with a probability *pGC*=0.70 [16]. Thus, the parameter *b_0_* (*b_0_* = 2 · *p_GC_* – 1) is about 30 times higher in humans than in yeast. Conversely, gene conversion tracts are on average about 4 times longer in yeast than in mammals [15,44]. Thus, for a same population-scaled recombination rate (*N_e_ r*), the intensity of gBGC would be about 7 times stronger in humans than in yeast. This example illustrates that because of variations in *L* and *b_0_*, the gBGC model does not necessarily predict a good correlation between population-scaled recombination rate and GC-content *across* species. In fact, to test the predictions of the gBGC model, it is more appropriate to investigate correlations between base composition and recombination rate *within* genomes, so that the other parameters (*N_e_*, *L* and *b_0_*) can be controlled for.

### The impact of gBGC on bacterial genome evolution is quantitatively strong

The observations presented previously are qualitatively consistent with the gBGC model. However, they do not provide a quantification of the impact of recombination on bacterial genomes: to what extent might this model account for the strong variations of GC-content observed across bacterial species? The gBGC model predicts that, all else being equal, the present-day GC-content of a genome should directly reflect its average recombination rate over long evolutionary time. To test this prediction, it is important to take into account two difficulties. First, recombination rates measured in extant populations reflect recent events (more recent than the coalescent time, i.e. of the order of *N_e_* generations), and hence may not correspond to the average recombination rate over times necessary for genomic GC-content to evolve significantly (i.e. inter-species divergence times). Second, the precision in the estimate depends on the physical scale at which recombination is measured. To illustrate these points let us consider the human genome, where the impact of gBGC is well documented [6]. At the gene scale, the correlation between present-day recombination rate (measured in a 10-kb window, centered on the middle of the gene, using HapMap genetic map [34]), and the gene GC-content (at third codon position) is significant but quite weak (R^2^=0.035, p<10^-10^). However, at 1Mb scale the correlation is much stronger (R^2^=0.15; [9]). Furthermore, when GC-content variations and recombination rates are measured over the same evolutionary time period, the correlation becomes very strong (1Mb scale: R^2^=0.64; [45]).

To test whether the impact of gBGC in bacteria was comparable to what is observed in mammals, we first focused on *Streptococcus pyogenes*, one of the species for which the signature of gBGC is strong (Fig. 1, see also Sup. Mat.). We computed the population-scaled recombination rate (*rho*) for each gene of the core genome, using ClonalOrigin [33]. The correlation between *rho* and the GC-content at third codon position of each gene (GC3) is higher than what is observed in humans (R^2^=0.087; p<10^-9^). This result is remarkable, given that recombination rates are measured here at the gene scale (typically about 1kb).

To go further, we binned the data set into 20 groups of genes according to their GC3, and we computed the correlation between the average GC3 and the average *rho* of each bin. Our reasoning is that by computing average values, we should get estimates of *rho* that are more robust to measurement noise and to possible temporal variations in recombination rates. Using this approach, we observed a strong correlation between the GC3 and recombination (R^2^=0.60; Fig. 3A). To investigate the amplitude of this relation in the other bacterial species studied here, we used P_REC_ (an index based on the proportion of genes in a bin that are detected as recombinant by PHI), which provides a good estimate of the average recombination rate in a bin, and is much easier to compute than ClonalOrigin’s *rho*. We observed a significant correlation for 11 of the 14 species, and these significant correlations were positive in all cases, with R^2^ values comprised between 0.24 and 0.68 (on average R^2^=0.43; Fig. 3B). Thus, in many bacteria, the average recombination rate in a bin is a good predictor of its average GC-content. We performed an analogous analysis in human genes using jackknife sampling of the dataset to scale it to the size of bacterial datasets. The average correlation observed in humans is R^2^=0.55 (Fig 3A). Hence, in many bacteria, the intensity of the relationships between GC-content and recombination is comparable to that observed in humans, where the impact of gBGC on base composition is known to be strong [45]. This is consistent with the hypothesis that on the long term, the gBGC process can have a major influence on the evolution of base composition in bacteria.

### A dynamic model of the evolution of nucleotide composition: long-term equilibrium vs.short-term disequilibrium

If the base composition of a genome is at evolutionary equilibrium then, by definition, the number of A/T→G/C substitutions must be equal to the number of G/C→A/T substitutions. Hildebrand and colleagues [23] noted that in a large majority of bacterial genomes (94/149), the number of G/C→A/T changes (inferred from the comparison of closely related organisms) exceeds the number of A/T→G/C changes. Given that genomic base composition strongly fluctuates over long evolutionary times (as demonstrated by the wide distribution of GC-content across bacterial species), it is not surprising that many genomes are not at equilibrium. However, what is unexpected is that this non-stationarity predominantly leads to loosing GC-content: *a priori*, at the scale of the entire bacterial biodiversity, one would expect to observe as many GC-increasing genomes as GC-decreasing genomes. One possible explanation is that the observed excess G/C→A/T changes among closely related genomes corresponds to polymorphic mutations, which eventually do not reach fixation because either selection or gBGC favors GC-alleles over AT-alleles [23]. Hildebrand and colleagues observed an excess of G/C→A/T changes even in bacterial genomes that show no evidence of recent recombination population-wise (i.e. *N_e_r* = 0) [23]. They therefore rejected the hypothesis that the fixation bias could be due to gBGC. However, this conclusion relies on one important assumption: that the *N_e_r* parameter measured in extant populations reflects the long-term average recombination rate. In fact it is expected that *N_e_* (and hence *N_e_r*) should fluctuate over time, as populations go through periods of bottlenecks and expansion. Immediately after a bottleneck, *N_e_r* would be close to 0, and hence genomes should accumulate G/C→A/T substitutions. However, on the long term, this can be compensated by an increase in GC-content when the effective population size becomes larger (and hence *B* > 1). Thus, the base composition of genomes may remain above the mutational equilibrium on the long term, even if many lineages go through periods during which *N_e_r* is null (and hence *B*=0). Interestingly, the rare species for which the long-term recombination rate is effectively null (typically endosymbiotic bacteria), generally have very AT-rich genomes [46], as predicted by the gBGC hypothesis.

### Universal mutational bias and selection on the strength of gBGC

Several lines of evidence suggest that the pattern of spontaneous mutations is biased towards AT nucleotides, in Eukaryotes as well as Prokaryotes [22,23,47]. Under such a bias, it is expected that the selective pressure to reduce mutation rate should generally favor a GC-biased DNA repair machinery. Recombination and repair are tightly linked processes that use many common pathways. In yeast, the analysis of conversion tracts in meiotic product indicates that the conversion bias is most probably due to the mismatch repair machinery (MMR) [48]. The MMR components involved in homologous recombination (MutS, MutL) are generally conserved between Bacteria and Eukaryotes. Hence, in Bacteria as well as Eukaryotes, gBGC could be the secondary effect of a selection for biased repair mechanisms. However, population genetics model show that when gBGC is strong, it drives the fixation of deleterious mutations [49]. Thus, one interesting hypothesis is that in highly recombining species, selection might favor unbiased repair mechanisms (i.e., values of *b_0_* close to 0 in equation (1)), so as to limit the deleterious consequences of strong gBGC. *H. pylori* is notorious for being highly recombining [50,51], as confirmed by our results. It is interesting that we have found no evidence for gBGC in this species, and recombining genes even seem to have slightly lower GC-content than non recombining genes. This trend is however relatively weak because there is no significant trend of variation of recombination rates among classes of homogeneous GC-content, as shown in fig. 3B. One possible explanation for the absence of gBGC in *H. pylori* could hence be that *b_0_* is null in this species, in compensation of a very high rate of recombination.

Although the MMR pathway is a good candidate as a molecular source of gBGC in Bacteria, the association of gBGC with MutSL genes is not straightforward. These genes are absent from three of our genome datasets, *C. jejuni, H. pylori* and *Bifidobacterium longum,* resulting from ancestral losses in Delta-Proteobacteria and Actinobacteridae, respectively [52]. In *H. pylori*, we indeed find no evidence of gBGC while the genomes are recombining at high frequency [50,51]. In *C. jejuni* and *B. longum*, however, we observe patterns similar to the other bacterial datasets that are in support of the existence of gBGC, indicating that it does not depend on the presence of a typical MutSL complex. The existence of gBGC in Bacteria and Eukaryotes however suggests that it may have been present in the last universal common ancestor of all cellular life forms (LUCA). Unfortunately, the only archaeal dataset matching our criteria was a group of *Sulfolobus* sp. genomes for which our analysis showed few evidence of recombination (Table 1), in agreement with the previously described isolation of endemic clades in this group [42].

We do not claim that gBGC is the unique determinant of base composition in bacterial genome: in fact there is evidence that mutation patterns vary significantly among species [22], and these variations are expected to contribute to differences in genome base composition. However, the model we propose provides a simple explanation for several important results of comparative bacterial genomics. First, gBGC can explain why bacterial genomes can maintain a high GC-content, even though the pattern of mutation is universally AT-biased [22,23]. Second, gBGC can explain some of the intragenomic heterogeneity in GC-content observed in bacterial genomes. Indeed, we observe that genes with evidence of recombination display on average substantially higher GC-content than other genes. This observation also suggests that the probability of recombination is variable among genes in the genome, as proposed under some speciation models [53]. Furthermore, given that recently acquired genes tend to be AT-rich, gBGC would contribute to their progressive enrichment in GC-content [54,55].

### gBGC, a new component of the neutral theory of evolution

The variations of GC-content in Bacteria have long remained unexplained. The results presented here highlight a strong relationship between the GC-content of genes and their history of recombination. This result, and the observation that bacterial genomes are generally above the GC-content predicted from their mutational bias towards AT, are fully consistent with the existence of gBGC. To explain our results under a selective model, one would have to hypothesize: i) that all bacterial species are under the same selection for higher GC throughout their genome, ii) that this selective pressure affects all positions of a genome, independently of gene function and expression, and iii) that the efficiency of selection varies with recombination (Hill-Robertson interference). Incidentally, if the correlations between GC-content and recombination (Figure 3B) were due to Hill-Robertson interference, this would imply that all regions in a genome (except possibly the most GC-rich) are mal-adapted. We favor the gBGC model because it is much more parsimonious, and it relies on mechanisms that have already been uncovered in Eukaryotes. Ultimately, it will be possible to test experimentally the existence of gBGC by analyzing recombination products in bacteria.

Our discovery is important because gBGC has been shown to interfere with the efficiency of selection in Eukaryotes, and to lead to false positives in the search for regions under positive selection in a genome. The prevalence – if not universality – of this phenomenon underlines the importance of incorporating gBGC in the set of evolutionary forces to be considered when searching for signature of adaption in genomes.

## Materials and Methods

### Genomic datasets

We used the HOGENOM database [27] to select sets of genome sequences comprising at least six closely related strains or species. The selection of closely related genomes was based on a genomic distance derived from the HOGENOM6 database. For each pair of genomes present in the database, we computed the best identity score, as obtained using BLAST, within each family of homologous proteins as defined in HOGENOM, and averaged these scores to obtain a global similarity score *s*. We then took 1-*s* as a distance and selected groups of genomes with at least 6 members and a distance lower than 0.15. This criterion left 21 groups of species representing a variety of bacterial and archeal species (Table 1). For each gene family, CDSs were extracted using ACNUC Python API [56] and re-aligned with MUSCLE [57] using default parameters. GC% and codon frequencies were computed using custom Python scripts.

### Detection of recombining genes

Detection of recombination based on multiple alignments is expected to be sensitive to both the number of sequences aligned, and the total length of the alignment. For this reason, we used only the universal unicopy genes of each species, and selected the 900 central positions of each nucleotide alignment (remaining positions as well as genes shorter than this threshold were ignored in our analysis). We used the software Phipack implementing the PHI test [28] (parameters: window size of 100bp, *p*-value computed from 1,000 permutations) to test if a gene family alignment contained evidence for recombination. Only families for which the site permutation test could be performed were considered, i.e. where the phylogenetic signal was sufficient to accept or reject the hypothesis of recombination. An alignment was determined to be “recombinant” if the *p*-value of the permutation test was lower than 0.05. To confirm the results obtained with PHI, we also used three other tests of recombination: NSS [58], MaxChi2 [59] and Geneconv [60]. NSS and MaxChi2 statistics were computed together with PHI (as implemented in Phipack package) using the same permutations to compute *p*-values. A consensus of PHI, NSS and MaxChi2 was done, classifying as recombinant or non-recombinant those gene families consistently detected by all those three methods as recombinant or non-recombinant, respectively; families with disagreeing results were discarded for the consensus analysis. Geneconv was run with the parameters “-GScale=1 -Numsims=10000 -Maxsimglobalpval=0.05”, and a gene family was considered recombinant when at least one significant global recombinant fragment was reported; other families were considered non-recombinant. As Geneconv differs from the other methods on the nature of the reported evidence, it was not considered when defining a consensus classification. Individual tests and their consensus yielded quite different classifications of gene families, but all led to qualitatively very similar results (see Fig. S1, S3 and S4). To test for the presence of gBGC, we performed Student’s *t*-tests comparing the mean GC-content (at all or at each separate codon position) of recombinant versus non-recombinant core gene family sets.

### Estimation of recombination rate

We used ClonalOrigin [33] to estimate the recombination rate on full-length core gene family alignments of 900bp or more. As ClonalOrigin inferences are highly demanding in computation time and power, we only performed this analysis on the moderately sized dataset of *S. pyogenes* genomes (*Spyo*). The program was used with default parameter weights, running the MCMC with 1,000,000 burn-in generations and 500,000 generations sampled every 1,000 (command line was: ‘warg -a 1,1,0.1,1,1,1,1,1,0,0,0 -x 1000000 -y 500000 -z 1000’). Families for which this task could not be finished in less than 1 month computation were discarded, yielding a total dataset of 478 gene families (out of 496 core ≥900bp-long ones). A number of genes also exhibited very high variance in their estimates of *rho* and their exclusion yielded better correlations with GC3.

### Recombination inference in intergenes

Intergenic regions are evolving fast, leading to inaccurate alignments that are not amenable to robust detection of recombination as performed on coding sequences. We thus chose to test the relationship of intergenic GC% with recombination in a gene-centered way: for each core protein-coding gene tested with PHI, we considered non-coding regions up to 400bp on both sides of a CDS and averaged their GC%, provided they were of a size larger than 50bp each (to avoid stochastic errors due to too small number of observed nucleotides). A measure of intergenic GC% per core gene family was obtained as the mean over all genomes of the previous values, and was associated to the recombinant/non-recombinant status of the gene family for further testing.

### Frequencies of optimal codons

Optimal codons for each amino acid were computed by comparing synonymous codon frequencies within CDSs encoding ribosomal proteins (as defined from HOGENOM family annotations) versus all other CDSs (RP method). Under the hypothesis of selection for highly expressed genes to be adapted to the tRNA pool, codons statistically enriched in ribosomal proteins (Chi-squared test based on a 2x2 contigency table of counts of occurrence at a focal codon against those of its synonyms in ribosomal vs. other protein-coding genes, with *p*-value <0.001) were considered as “optimal”; others were classified as “non-optimal”. (Table S2). We then computed the absolute frequency of optimal (Fop) and non-optimal codons (Fnop) over all coding sequences. Fop and Fnop were calculated separately for codons pooled by composition at the third position, i.e. ending in A/U or G/C. As there is a debate on whether this method is appropriate to define optimal codons [30–32], we also used an alternative definition and took optimal codons datasets from a previous exhaustive survey of Hershberg and Petrov [30] (HCB method). A set of optimal codons was selected when determined in Hershberg and Petrov [30] study for several strains with a strong consensus, i.e. when >60% documented strains agreed on the preferred codon and remaining strains preferred a codon with the same composition (A/U or G/C) at the third position (found in all datasets but *A. baumanii*) (Table S3).

## Financial Disclosure

This work was supported by French National Research Agency (ANR) grant Ancestrome (ANR-10-BINF-01-01) and EcoGenome (BLAN-08-0090). (http://www.agence-nationale-recherche.fr/) FL received a doctoral scholarship from Ecole Normale Supérieure de Lyon (http://www.ens-lyon.eu/) and is supported by ERC grant BIG_IDEA 260801 (http://erc.europa.eu).

The funders had no role in study design, data collection and analysis, decision to publish, or preparation of the manuscript.

## Acknowledgements

We thank Gergely Szöllősi, Paul Sharp and Eduardo PC Rocha for their thoughtful comments and advice about controls to include in the present work. We thank Simon Penel and Vincent Miele for providing scripts and data used to process the genomic data from HOGENOM database.

## Supplementary figures

**Figure S1 – Comparison of several recombination detection programs on the inferred effect of recombination on core genes GC-content**

Legend as in Fig. 1.

**Figure S2 – Difference of GC% between intergenes located next to recombining vs. non−recombining genes**

Difference in average GC-content of intergenic regions around each single-copy core gene which had a conclusive result of the PHI test. Individual intergenic GC% values were computed as the average of both flanking intergenes when they were 50bp or longer, measured on at most 400bp away of the reference gene. Intergenes were classified as recombinant and non-recombinant as was the neighbouring gene based on the PHI test. A positive difference indicates that intergenes next to recombinant families are enriched in GC. Symbols and dataset abbreviations as in Fig. 1.

**Figure S3 – Comparison of several recombination detection programs on the inferred effect of recombination on codon usage of core genes, based on RP optimal codons.**

Legend as in Fig. 2.

**Figure S4 – Comparison of several recombination detection programs on the inferred effect of recombination on codon usage of core genes, based on HCB optimal codons from Hershberg and Petrov (2009).**

Legend as in Fig. 2.

## Supplementary Tables

**Table S1 – Detailed results of statistical tests of difference of GC% and Fop/Fnop between recombining and nonrecombining core genes, varying recombination detection methods and tested alignment length.**

**Table S2 – Sets of optimal/non-optimal codons defined using RP method**

**Table S3 – Sets of optimal/non-optimal codons defined using HCB method**

Data derived from dataset established by Hershberg and Petrov[30].

## References

1. Doolittle WF (2013) Is junk DNA bunk? A critique of ENCODE. Proc Natl Acad Sci 110: 5294–5300. doi:10.1073/pnas.1221376110.

2. Bernardi G, Olofsson B, Filipski J, Zerial M, Salinas J, et al. (1985) The mosaic genome of warm-blooded vertebrates. Science 228: 953–958.

3. Sueoka N (1962) ON THE GENETIC BASIS OF VARIATION AND HETEROGENEITY OF DNA BASE COMPOSITION. Proc Natl Acad Sci 48: 582–592.

4. McCutcheon JP, Moran NA (2010) Functional convergence in reduced genomes of bacterial symbionts spanning 200 My of evolution. Genome Biol Evol 2: 708–718. doi:10.1093/gbe/evq055.

5. Pagani I, Liolios K, Jansson J, Chen I-MA, Smirnova T, et al. (2012) The Genomes OnLine Database (GOLD) v.4: status of genomic and metagenomic projects and their associated metadata. Nucleic Acids Res 40: D571–D579. doi:10.1093/nar/gkr1100.

6. Duret L, Galtier N (2009) Biased gene conversion and the evolution of mammalian genomic landscapes. Annu Rev Genomics Hum Genet 10: 285–311. doi:10.1146/annurev-genom-082908-150001.

7. Ratnakumar A, Mousset S, Glémin S, Berglund J, Galtier N, et al. (2010) Detecting positive selection within genomes: the problem of biased gene conversion. Philos Trans R Soc Lond B Biol Sci 365: 2571–2580. doi:10.1098/rstb.2010.0007.

8. Spencer CC, Deloukas P, Hunt S, Mullikin J, Myers S, et al. (2006). The influence of recombination on human genetic diversity. PLoS Genetics, 2:e148. doi:10.1371/journal.pgen.0020148.

9. Duret L, Arndt PF (2008) The Impact of Recombination on Nucleotide Substitutions in the Human Genome. PLoS Genet 4: e1000071. doi:10.1371/journal.pgen.1000071.

10. Webster MT, Hurst LD (2012) Direct and indirect consequences of meiotic recombination: implications for genome evolution. Trends Genet 28: 101–109. doi:10.1016/j.tig.2011.11.002.

11. Nagylaki T (1983) Evolution of a finite population under gene conversion. Proc Natl Acad Sci U S A 80: 6278–6281.

12. Galtier N, Duret L (2007) Adaptation or biased gene conversion? Extending the null hypothesis of molecular evolution. Trends Genet TIG 23: 273–277. doi:10.1016/j.tig.2007.03.011.

13. Galtier N, Duret L, Glémin S, Ranwez V (2009) GC-biased gene conversion promotes the fixation of deleterious amino acid changes in primates. Trends Genet TIG 25: 1–5. doi:10.1016/j.tig.2008.10.011.

14. Necşulea A, Popa A, Cooper DN, Stenson PD, Mouchiroud D, et al. (2011) Meiotic recombination favors the spreading of deleterious mutations in human populations. Hum Mutat 32: 198–206. doi:10.1002/humu.21407.

15. Mancera E, Bourgon R, Brozzi A, Huber W, Steinmetz LM (2008) High-resolution mapping of meiotic crossovers and non-crossovers in yeast. Nature 454: 479–485. doi:10.1038/nature07135.

16. Williams A, Geneovese G, Dyer T, Truax K, Jun G, et al. (2014) Non-crossover gene conversions show strong GC bias and unexpected clustering in humans. bioRxiv: 009175. doi:10.1101/009175.

17. Capra JA, Pollard KS (2011) Substitution patterns are GC-biased in divergent sequences across the metazoans. Genome Biol Evol 3: 516–527. doi:10.1093/gbe/evr051.

18. Escobar JS, Glémin S, Galtier N (2011) GC-biased gene conversion impacts ribosomal DNA evolution in vertebrates, angiosperms, and other eukaryotes. Mol Biol Evol 28: 2561–2575. doi:10.1093/molbev/msr079.

19. Pessia E, Popa A, Mousset S, Rezvoy C, Duret L, et al. (2012) Evidence for widespread GC-biased gene conversion in eukaryotes. Genome Biol Evol 4: 675–682. doi:10.1093/gbe/evs052.

20. Foerstner KU, von Mering C, Hooper SD, Bork P (2005) Environments shape the nucleotide composition of genomes. EMBO Rep 6: 1208–1213. doi:10.1038/sj.embor.7400538.

21. Sueoka N (1988) Directional mutation pressure and neutral molecular evolution. Proc Natl Acad Sci U S A 85:2653.

22. Hershberg R, Petrov DA (2010) Evidence That Mutation Is Universally Biased towards AT in Bacteria. PLoS Genet 6: e1001115. doi:10.1371/journal.pgen.1001115.

23. Hildebrand F, Meyer A, Eyre-Walker A (2010) Evidence of Selection upon Genomic GC-Content in Bacteria. PLoS Genet 6. doi:10.1371/journal.pgen.1001107.

24. Touchon M, Hoede C, Tenaillon O, Barbe V, Baeriswyl S, et al. (2009) Organised genome dynamics in the Escherichia coli species results in highly diverse adaptive paths. PLoS Genet 5: e1000344. doi:10.1371/journal.pgen.1000344.

25. Rocha EPC, Feil EJ (2010) Mutational Patterns Cannot Explain Genome Composition: Are There Any Neutral Sites in the Genomes of Bacteria? PLoS Genet 6: e1001104. doi:10.1371/journal.pgen.1001104.

26. Raghavan R, Kelkar YD, Ochman H (2012) A selective force favoring increased G+C content in bacterial genes. Proc Natl Acad Sci 109: 14504–14507. doi:10.1073/pnas.1205683109.

27. Penel S, Arigon A-M, Dufayard J-F, Sertier A-S, Daubin V, et al. (2009) Databases of homologous gene families for comparative genomics. BMC Bioinformatics 10: S3. doi:10.1186/1471-2105-10-S6-S3.

28. Bruen TC, Philippe H, Bryant D (2006) A Simple and Robust Statistical Test for Detecting the Presence of Recombination. Genetics 172: 2665–2681. doi:10.1534/genetics.105.048975.

29. McVean GA, Charlesworth B (2000) The effects of Hill-Robertson interference between weakly selected mutations on patterns of molecular evolution and variation. Genetics 155: 929–944.

30. Hershberg R, Petrov DA (2009) General Rules for Optimal Codon Choice. PLoS Genet 5: e1000556. doi:10.1371/journal.pgen.1000556.

31. Wang B, Shao Z-Q, Xu Y, Liu J, Liu Y, et al. (2011) Optimal Codon Identities in Bacteria: Implications from the Conflicting Results of Two Different Methods. PLoS ONE 6: e22714. doi:10.1371/journal.pone.0022714.

32. Hershberg R, Petrov DA (2012) On the Limitations of Using Ribosomal Genes as References for the Study of Codon Usage: A Rebuttal. PLoS ONE 7: e49060. doi:10.1371/journal.pone.0049060.

33. Didelot X, Lawson D, Darling A, Falush D (2010) Inference of Homologous Recombination in Bacteria Using Whole Genome Sequences. Genetics 186: 1435–1449. doi:10.1534/genetics.110.120121.

34. Frazer KA, Ballinger DG, Cox DR, Hinds DA, Stuve LL, et al. (2007) A second generation human haplotype map of over 3.1 million SNPs. Nature 449: 851–861. doi:10.1038/nature06258.

35. Ussery DW, Kiil K, Lagesen K, Sicheritz-Pontén T, Bohlin J, et al. (2009) The genus burkholderia: analysis of 56 genomic sequences. Genome Dyn 6: 140–157. doi:10.1159/000235768.

36. Joseph SJ, Didelot X, Gandhi K, Dean D, Read TD (2011) Interplay of recombination and selection in the genomes of Chlamydia trachomatis. Biol Direct 6: 28. doi:10.1186/1745-6150-6-28.

37. Achtman M, Zurth K, Morelli G, Torrea G, Guiyoule A, et al. (1999) Yersinia pestis, the cause of plague, is a recently emerged clone of Yersinia pseudotuberculosis. Proc Natl Acad Sci 96: 14043–14048. doi:10.1073/pnas.96.24.14043.

38. Keim P, Johansson A, Wagner DM (2007) Molecular Epidemiology, Evolution, and Ecology of Francisella. Ann N Y Acad Sci 1105: 30–66. doi:10.1196/annals.1409.011.

39. Supply P, Marceau M, Mangenot S, Roche D, Rouanet C, et al. (2013) Genomic analysis of smooth tubercle bacilli provides insights into ancestry and pathoadaptation of Mycobacterium tuberculosis. Nat Genet 45: 172–179. doi:10.1038/ng.2517.

40. Wattam AR, Williams KP, Snyder EE, Almeida NF Jr, Shukla M, et al. (2009) Analysis of ten Brucella genomes reveals evidence for horizontal gene transfer despite a preferred intracellular lifestyle. J Bacteriol 191: 3569–3579. doi:10.1128/JB.01767-08.

41. Whitaker RJ, Grogan DW, Taylor JW (2003) Geographic Barriers Isolate Endemic Populations of Hyperthermophilic Archaea. Science 301: 976–978. doi:10.1126/science.1086909.

42. Reno ML, Held NL, Fields CJ, Burke PV, Whitaker RJ (2009) Biogeography of the Sulfolobus islandicus pangenome. Proc Natl Acad Sci 106: 8605–8610. doi:10.1073/pnas.0808945106.

43. Balbi KJ, Rocha EPC, Feil EJ (2009) The Temporal Dynamics of Slightly Deleterious Mutations in Escherichia coli and Shigella spp. Mol Biol Evol 26: 345–355. doi:10.1093/molbev/msn252.

44. Cole F, Baudat F, Grey C, Keeney S, de Massy B, et al. (2014) Mouse tetrad analysis provides insights into recombination mechanisms and hotspot evolutionary dynamics. Nat Genet 46: 1072–1080. doi:10.1038/ng.3068.

45. Munch K, Mailund T, Dutheil JY, Schierup MH (2014) A fine-scale recombination map of the human–chimpanzee ancestor reveals faster change in humans than in chimpanzees and a strong impact of GC-biased gene conversion. Genome Res 24: 467–474. doi:10.1101/gr.158469.113.

46. Moran NA (2002) Microbial minimalism: genome reduction in bacterial pathogens. Cell 108: 583–586.

47. Lynch M (2010) Rate, molecular spectrum, and consequences of human mutation. Proc Natl Acad Sci 107: 961–968. doi:10.1073/pnas.0912629107.

48. Lesecque Y, Mouchiroud D, Duret L (2013) GC-Biased Gene Conversion in Yeast Is Specifically Associated with Crossovers: Molecular Mechanisms and Evolutionary Significance. Mol Biol Evol 30: 1409–1419. doi:10.1093/molbev/mst056.

49. Glémin S (2010) Surprising fitness consequences of GC-biased gene conversion: I. Mutation load and inbreeding depression. Genetics 185: 939–959. doi:10.1534/genetics.110.116368.

50. Suerbaum S, Smith JM, Bapumia K, Morelli G, Smith NH, et al. (1998) Free recombination within Helicobacter pylori. Proc Natl Acad Sci 95: 12619–12624. doi:10.1073/pnas.95.21.12619.

51. Falush D, Kraft C, Taylor NS, Correa P, Fox JG, et al. (2001) Recombination and mutation during long-term gastric colonization by Helicobacter pylori: Estimates of clock rates, recombination size, and minimal age. Proc Natl Acad Sci 98: 15056–15061. doi:10.1073/pnas.251396098.

52. Lin Z, Nei M, Ma H (2007) The origins and early evolution of DNA mismatch repair genes—multiple horizontal gene transfers and co-evolution. Nucleic Acids Res 35: 7591–7603. doi:10.1093/nar/gkm921.

53. Retchless AC, Lawrence JG (2007) Temporal fragmentation of speciation in bacteria. Science 317: 1093–1096. doi:10.1126/science.1144876.

54. Daubin V, Lerat E, Perrière G (2003) The source of laterally transferred genes in bacterial genomes. Genome Biol 4: R57. doi:10.1186/gb-2003-4-9-r57.

55. Daubin V, Ochman H (2004) Bacterial Genomes as New Gene Homes: The Genealogy of ORFans in E. coli. Genome Res 14: 1036–1042. doi:10.1101/gr.2231904.

56. Gouy M, Gautier C, Attimonelli M, Lanave C, di Paola G (1985) ACNUC--a portable retrieval system for nucleic acid sequence databases: logical and physical designs and usage. Comput Appl Biosci CABIOS 1: 167–172.

57. Edgar RC (2004) MUSCLE: a multiple sequence alignment method with reduced time and space complexity. BMC Bioinformatics 5: 113. doi:10.1186/1471-2105-5-113.

58. Jakobsen IB, Easteal S (1996) A program for calculating and displaying compatibility matrices as an aid in determining reticulate evolution in molecular sequences. Comput Appl Biosci CABIOS 12: 291–295.

59. Smith JM (1992) Analyzing the mosaic structure of genes. J Mol Evol 34: 126–129.

60. Sawyer S (1989) Statistical tests for detecting gene conversion. Mol Biol Evol 6: 526–538.

